# The interaction between the tumour suppressor Dlg1 and the MAGUK protein CASK is required for oriented cell division in mammalian epithelia

**DOI:** 10.1101/482588

**Authors:** Andrew P. Porter, Gavin White, Natalie A. Mack, Angeliki Malliri

## Abstract

Oriented cell divisions are important for the formation of normal epithelial structures. Dlg1, a tumour suppressor, is required for oriented cell division in *Drosophila* epithelia and chick neuroepithelia, but how Dlg1 is localised to the membrane and its importance in mammalian epithelia are unknown. Here we show that Dlg1 is required in non-transformed mammalian epithelial cells for oriented cell divisions, and for normal lumen formation in 3D culture. We demonstrate that CASK, a membrane-associated scaffold, is the factor responsible for Dlg1 membrane localisation during spindle orientation, and thereby identify a new cellular function for CASK. We show that depletion of CASK leads to misoriented divisions in 3D, and to the formation of multilumen structures in cultured kidney and breast epithelial cells. Blocking the direct interaction between CASK and Dlg1 with an interfering peptide disrupts spindle orientation and causes multilumen formation. We further show that the Dlg1-CASK interaction is important for the membrane localisation of the canonical LGN-NuMA complex, required for attachment of the mitotic spindle to the membrane and its correct positioning, as well as for astral microtubule stability. Together these results establish the importance of the CASK-Dlg1 interaction in oriented cell division and epithelial integrity.

## Introduction

Control of the orientation of cell division through regulating the orientation of the mitotic spindle plays an important role in developing and maintaining tissue architecture in both embryonic and adult tissues. Oriented divisions have been observed in a diverse range of organisms and cell types, including yeast, the fruit fly *Drosophila*, chick neuroepithelial and mammalian epithelial systems, and occur in both stem cells and differentiated tissues [1]. Regulated changes in spindle orientation can control the balance between proliferation and differentiation in stratified epithelia [2]. The role of spindle orientation in symmetrically dividing epithelial cells is less well understood. Epithelial cells tend to divide in the plane of the epithelium, which is important to integrate daughter cells within the epithelium and thereby maintain barrier function [2] and these oriented divisions are hypothesised to have a tumour suppressive function [3], although direct evidence of its importance for tumourigenesis is limited at this point.

A conserved set of proteins has been identified which regulate spindle orientation across a range of organisms and tissue types by defining membrane domains for the attachment of astral microtubules, which in turn orient the mitotic spindle [1]. These include Gαi, which is involved in the localisation of LGN (Leucine–Glycine–Asparagine) and subsequently NuMA (Nuclear and Mitotic Apparatus) to the plasma membrane (reviewed in [4]). In turn, NuMA binds to and stabilises astral microtubules, and recruits the microtubule-binding motor protein dynein [5]. In mammalian cells, pulling forces at the cell cortex are transmitted along astral microtubules which then act to correctly position the mitotic spindle.

In planar divisions of HeLa cells on L-shaped micropatterns, and in chick neuroepithelial cells, LGN localisation is regulated in part by interaction with Dlg (Discs Large); LGN membrane localisation is reduced following Dlg1 depletion [6], while in *Drosophila* follicular epithelia Dlg loss leads to redistribution of Pins (the *Drosophila* orthologue of LGN) [7]. However, in other systems, interaction with E-cadherin is required for localisation of LGN [8]. Whether Dlg1 plays a role in orienting the mitotic spindle along the apical-basal axis in non-transformed mammalian epithelial cells has not been determined, and the factor regulating Dlg1 membrane localisation in the context of spindle orientation has yet to be identified [9].

In this report we show that Dlg1 is required for spindle orientation in 3D cultures of untransformed mammalian epithelial cells, and identify the MAGUK protein CASK as the protein responsible for Dlg1 membrane localisation in the context of spindle orientation. By blocking CASK-Dlg1 binding we show that this protein-protein interaction is required for Dlg1 localisation, and subsequently the localisation of the LGN-NuMA complex, which binds the astral microtubules that ultimately orient the mitotic spindle. We also show that blocking the CASK-Dlg1 interaction leads to the formation of multilumen structures.

## Results and Discussion

### Dlg1 regulates spindle orientation and epithelial lumen formation in mammalian cells

MDCKII cells seeded onto Matrigel have the capacity to grow as cysts, reminiscent of those found in the mammalian kidney, with a hollow lumen surrounded by a single layer of epithelial cells. We knocked down Dlg1 using two independent siRNAs (Figure 1a) and saw that this disrupted normal lumen formation in 3D culture, giving rise to cysts with multiple lumens, as marked by strong apical actin staining (Figure 1b and 1c). Similarly, MDCKII cells grown for 8 to 10 days embedded in a Collagen I matrix produce cysts with a single lumen, as marked with apical actin and GP135/Podocalyxin staining (Figure 1d, top left panel). Cysts constitutively expressing an shRNA hairpin against Dlg1 displayed disrupted lumen development, with many cysts containing multiple lumens (Figure 1d and 1e). Staining for Dlg1 in these cysts revealed strong basolateral localisation of Dlg1 in control cysts, which is lost following Dlg1 knockdown (Figure 1d).

**Figure 1-.**
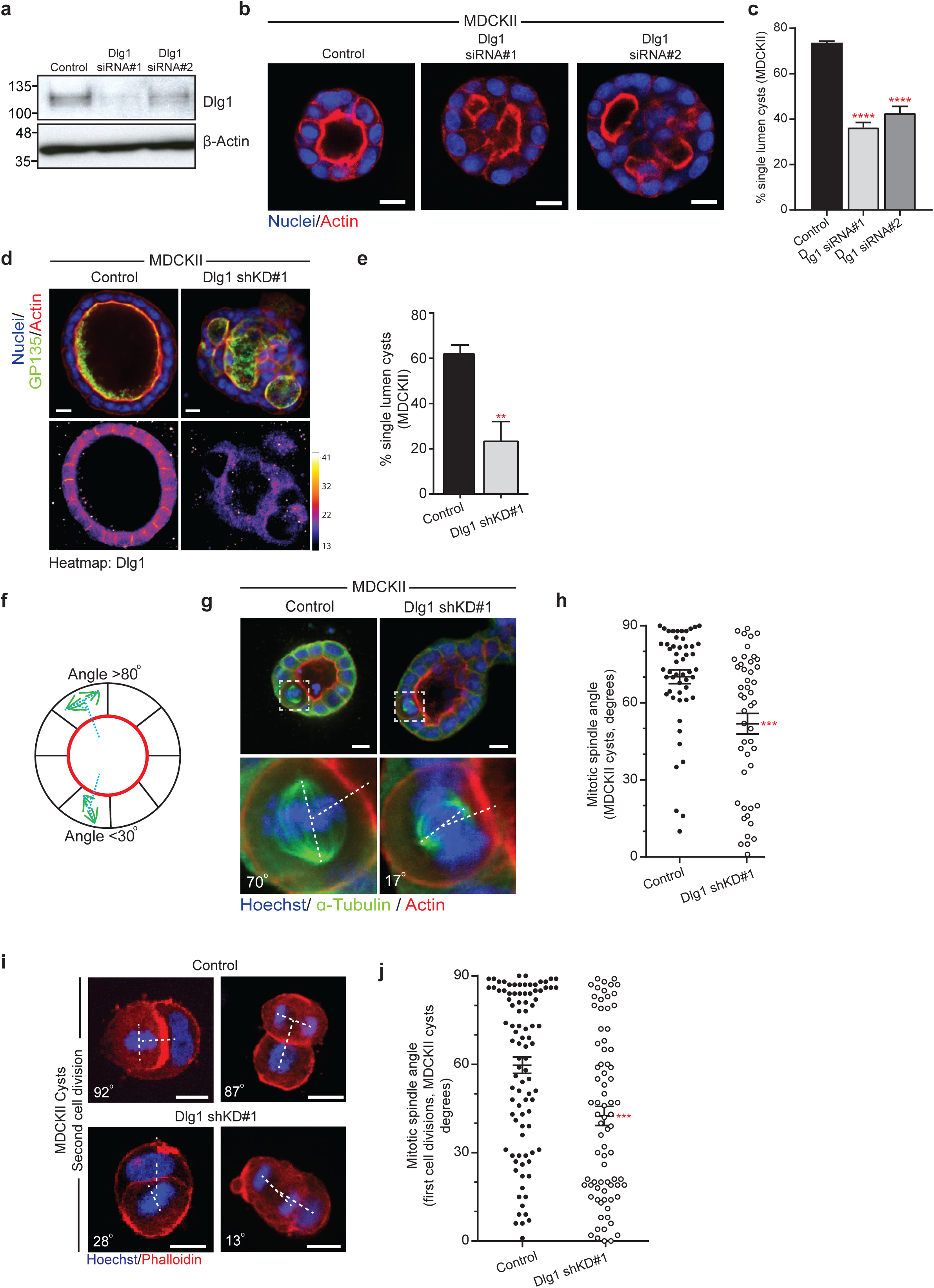
Dlg1 regulates epithelial lumen formation and mitotic spindle orientation. a) Western blot showing depletion of Dlg1 with two siRNAs. b) Confocal images of Control (Non- Targeting) or Dlg1 siKD#1 and siKD#2 cysts grown in 2% Matrigel, showing multilumen structures following Dlg1 depletion. c) Quantification of single lumen cysts from three independent Dlg1 knockdown experiments, n>40 cysts per experiment; **** p<0.0001 (One-way ANOVA.) d) Confocal images of control (Non-Targeting) or Dlg1 shKD#1 MDCKII cysts grown in Collagen I, stained for the indicated proteins to show lumens (upper panels) or Dlg1 levels (lower panels). e) Quantification of single lumen cysts from three independent Dlg1 knockdown experiments as in (d), n>100 cysts per experiment; ** p=0.0083. (unpaired Student’s t-test, two tailed). f) Schematic showing measurement of spindle orientation in MDCKII cysts showing a normally oriented division (>80°) and a misoriented division (<30°). g) Representative confocal images of MDCKII cysts grown in Collagen I and expressing Non-Targeting (Control) or Dlg1 shRNA. Mitotic cells are magnified in inserts (lower panels) and annotated to show spindle angles. h) Quantification of spindle angles in constitutive knockdown experiments as in (g), n=52/47 cells pooled from three independent experiments; *** p=0.0004. Data analysed using Mann-Whitney test. i) Images of early cell divisions (metaphase, left panel, and telophase, right panel) in developing MDCKII cysts grown in Collagen I. Shown are representative images from Control (Non-Targeting) and Dlg1 constitutive knockdown cells, showing misoriented divisions following Dlg1 knockdown. j) Quantification of early cell divisions in MDCKII cysts from experiments as in (i), n=95/78 from three independent experiments; *** p<0.0001 (Mann-Whitney test). Error bars show mean +/- SEM. All scale bars are 10 µm.

Normal lumen formation in MDCKII cysts has been linked to tight regulation of the mitotic spindle [10, 11], so we investigated whether there was a disruption to the orientation of the mitotic spindle in MDCKII cells depleted for Dlg1. We measured the angle of divisions in mitotic cells relative to the apical surface of the dividing cell (Figure 1f). In control cells expressing a non-targeting shRNA the mitotic spindle tended to align orthogonally to the apical surface of the dividing cell, whereas mitotic divisions in Dlg1 knockdown cells were much more widely distributed and essentially randomised (Figure 1g and 1h). This demonstrates a requirement for Dlg1 in spindle orientation in non-transformed mammalian epithelial cells.

Dlg1 is involved in the establishment of polarity in fruit flies [12] and required for adherens junction formation in *C. elegans* [13], and while its role in mammalian epithelial polarity is less clear, if loss of Dlg1 globally affected the polarity of the cyst this might indirectly affect spindle orientation. We therefore investigated spindle orientation in 2D cultures of confluent MDCKII cells, where cells have a strong extrinsic polarity signal from their attachment to the glass coverslip. Control cells aligned their mitotic spindles tightly to the plane of the coverslip, whereas we observed a significant tilting of cell divisions following Dlg1 knockdown (Supplementary Figure 1a, quantified in 1b). Dlg1 localises to lateral cell contacts and therefore loss of Dlg1 may affect spindle orientation through a general defect in cell-cell adhesion. To exclude an indirect effect of Dlg1 via reduced adhesion to adjacent cells we seeded single cells in collagen and measured the orientation of the second division in 3D where cells have only one, apical neighbour (Figure 1i). In control cells, the mitotic division tended to be orthogonal to the adjacent, apical cell (example image of metaphase and telophase cells in Figure 1i, quantified in 1j). Upon knockdown of Dlg1, a randomisation of the angle of cell division was observed (Figure 1i, 1j), with many cells dividing directly towards the adjacent cell, indicating that Dlg1 is required for orientation of the mitotic spindle at least in part by cell-autonomous mechanisms independent of lateral cell-cell adhesion.

### CASK is required for Dlg1 membrane localisation, lumen formation and spindle orientation

We next set out to investigate how Dlg1 is localised to the lateral membranes. Dlg1 has not been reported to bind directly to cell membranes, and does not contain a membrane localisation domain; instead another factor must be involved in its recruitment to the membrane. It has been reported that the MAGUK protein CASK is involved in Dlg1 localisation to the plasma membrane in some tissues [14, 15]. CASK is a multi-domain scaffolding protein with unusual magnesium-independent kinase activity [16], and contains a Hook domain, which mediates interaction with the cytoskeleton and allows for membrane binding [14]. Like Dlg1, deletion of CASK in mice is lethal [17], while mutations in CASK are associated with X-linked mental dysfunction [18] and craniofacial abnormalities [18] in humans. It is well-characterised in neurons [19] where it regulates both trafficking [20] and transcriptional pathways [21]; however, its function in epithelial cells is not understood. We stained 2D cultures of MDCKII cells and saw that CASK colocalised with Dlg1 at cell membranes in interphase and mitosis (Figure 2a). To test whether CASK was important for Dlg1 localisation we generated MDCKII cells with two different shRNAs against CASK under control of a doxycycline-inducible promoter. Treatment of cells with doxycycline efficiently depleted CASK (Figure 2b). We grew these cells in 3D culture and stained for Dlg1. In contrast to the clear basolateral staining of Dlg1 seen in the control (minus dox) cells, basolateral Dlg1 staining is completely lost following CASK knockdown (plus dox) (Figure 2c). (Of note, we have not been able to successfully image CASK in 3D cultures due to poor antibody staining in this context.) Interestingly, other basolateral markers, such as β-Catenin, still show a clear basolateral membrane staining in CASK knockdown cysts (Supplementary Figure 2a), and individual cells exhibit clear apical-basal polarity with strong apical GP135/Podocalyxin staining (Figure 2d), indicating that loss of CASK does not lead to a general loss of cell polarity which might indirectly account for the loss of Dlg1 localisation.

We tested whether CASK and Dlg1 interact in MDCKII cells, and were able to co-immunoprecipiate CASK and Dlg1 (Supplementary Figure 2b). Given that CASK depletion leads to a loss of Dlg1 localisation, we reasoned that CASK depletion alone might be sufficient to disrupt normal lumen formation. We assessed lumen formation in 3D culture using our inducible knockdown cells, and saw that treatment of cells with doxycycline led to a significant decrease in normal cysts, with many more cysts containing multiple lumens compared with untreated cells (Figure 2d and 2e). To demonstrate that CASK depletion itself was responsible for the formation of multiple lumen cysts, we generated a rescue system. First we generated MDCKII cells with constitutive expression of an shRNA against CASK, which lead to depletion of CASK protein. We confirmed that these cells also demonstrated the multiple lumen phenotype (Supplementary Figure 2c, 2d; using the same control as in Figure 1e). Next, we used the pRetro-X-Tight dual plasmid system to allow doxycycline inducible expression of full-length shRNA-resistant CASK. Addition of doxycycline efficiently restores CASK protein expression in these cells (Figure 2f). In the absence of doxycycline, cells grown in 3D culture produce multilumen structures (Figure 2g, third panel). Upon addition of doxycycline, CASK expression is restored, and these cells show a marked decrease in the number of multilumen cysts formed (Figure 2g, fourth panel, quantified in 2h). We were able to observe a recovery of Dlg1 membrane localisation in our rescue cells, indicating that CASK itself specifically regulates Dlg1 localisation (Supplementary Figure 2e).

**Figure 2-.**
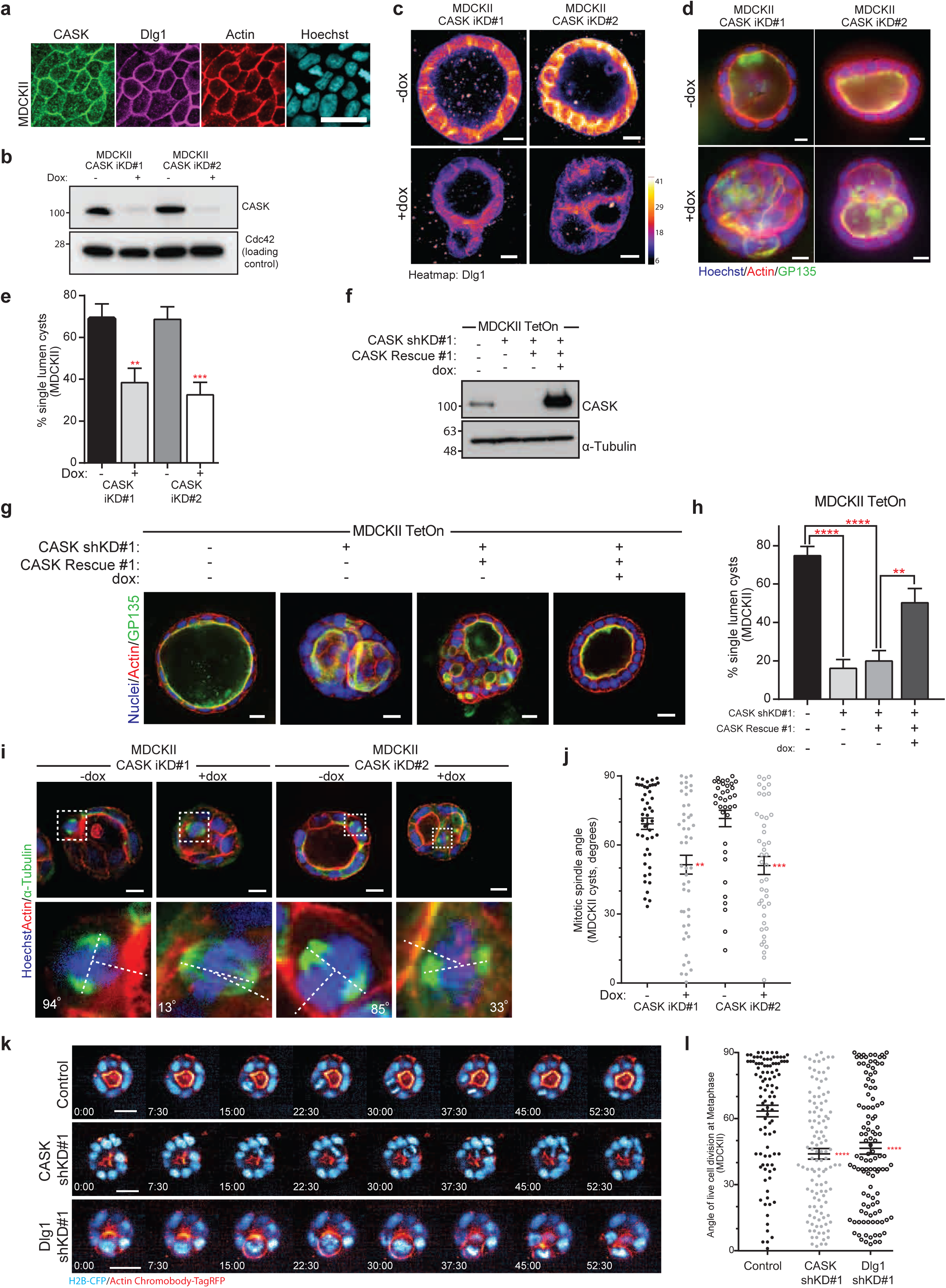
The Dlg1 interactor CASK is required for normal lumen formation and spindle orientation. a) Widefield images showing CASK and Dlg1 colocalisation in MDCKII cells. Scale bar 30 µm. b) Western blot of MDCKII cell lysates from two shRNA inducible knockdown lines (CASK iKD#1 and CASK iKD#2) showing reduction of CASK protein levels following shRNA expression upon treatment with doxycycline (+dox). c) Confocal images of MDCKII cysts grown in Collagen I and stained for Dlg1 protein, showing basolateral localisation in control cysts (-dox), and a reduction following CASK knockdown (+dox). d) Widefield images of MDCKII cysts grown in Collagen I show a multilumen phenotype following CASK knockdown upon addition of doxycycline. e) Quantification of single lumen cysts from experiments as in (d). More than 100 cysts examined per experiment. Total number of independent experiments was 6/6/8/8 respectively; ** p=0.0059 ***, p=0.0004 (unpaired t-test, 2 tailed). f) Western blot of MDCKII cells showing constitutive knockdown of CASK and re-expression of shRNA-resistant wild-type CASK following addition of dox (+dox) restoring CASK expression levels. g) Confocal images of MDCKII cysts of (f) showing multilumen formation in the absence of CASK (-dox), which is rescued by re-expression of CASK (+dox), right panel. h) Quantification of single lumen cysts from experiments as in (g) with more than 100 cysts examined per experiment. Total number of independent experiments was 6/3/5/5 respectively; **** p=2.1×10^−5^ for CASK shKD#1 and 7.4×10^−6^ for CASK shKD#1 + CASK rescue –dox, ** p=0.0049 (one-way ANOVA). i) Confocal images of dividing cells in control (-dox) and CASK inducible knockdown cysts (+dox) grown in Collagen I; insets show individual divisions annotated to demonstrate how spindle orientation is measured. j) Quantification of spindle orientations in control (-dox) and CASK inducible knockdown (+dox) cysts as in (i). n=47/48/35/43 cells pooled from three independent experiments; ** p=0.0025 and *** p=0.0004 for CASK iKD#1 (+dox) and CASK iKD#2 (+ dox) respectively (Mann-Whitney test). k) Representative images from movies of dividing cells within MDCKII cysts, grown on Matrigel, showing a normal division (top), and misoriented divisions (middle, CASK shKD#1 and bottom, Dlg1 shKD#1). Time shown in minutes. l) Quantification of cell division angles from live imaging experiments, only from cells in metaphase (n=98/117/112 cells from three independent replicates); **** p=8.2×10^−6^ and 4.0×10^−5^ (Kruskal-Wallis test) for CASK shKD#1 and Dlg1 shKD#1 respectively. Error bars show mean +/- SEM. All scale bars are 10 µm except where stated.

CASK is a highly conserved protein, so we wanted to test whether CASK depletion would have similar effects on lumen formation in other mammalian epithelial cell types. To this end we made use of the normal breast epithelial cell line MCF10A, which forms acini when grown in Matrigel, with a single hollow lumen surrounded by a layer of epithelial cells. These are similar to the acini found in normal human breast tissue. We infected MCF10A cells with a retroviral plasmid containing a constitutively expressed shRNA against CASK, or a scrambled control, and grew the cells in 3D culture. After 10 days of growth, the control cells predominantly formed normal acini with a single lumen (Supplementary Figure 2f), whereas CASK knockdown MCF10A formed acini with multiple lumens (Supplementary Figure 2f, quantified in Supplementary Figure 2g). This demonstrates that CASK is important for lumen formation in non-cancerous human breast epithelial cells.

Next we tested whether CASK itself was required for normal spindle orientation. We used our inducible CASK knockdown MDCKII cells and measured mitotic spindle orientation in cysts grown in 3D. The addition of doxycycline, which leads to knockdown of CASK, induced a change in spindle orientation compared with control (minus doxycycline) cysts, with the spindle orientation becoming essentially randomised following CASK depletion (Figure 2i and 2j). We also observed this effect in 2D culture where we saw that the mitotic spindles were tilted away from the plane of the coverslip following constitutive CASK knockdown, similarly to the tilting observed with Dlg1 knockdown (Supplementary Figure 2h, 2i). Next we determined if this role for CASK in spindle orientation was conserved in human epithelial cells. We measured the angle of the mitotic spindle in MCF10A cells grown on glass coverslips; in scrambled control cells the mitotic spindle clearly aligned with the plane of the coverslip, whereas in CASK knockdown cells the mitotic spindles became tilted (Supplementary Figure 2j and 2k). These experiments demonstrate a new role for CASK in epithelial cells, in maintaining epithelial integrity and the orientation of cell division.

Cell division is a dynamic process, the details of which may not be fully captured by snapshots in fixed cells. We therefore developed a method for assaying spindle orientation in live cysts. We transfected MDCKII cells with a nuclear marker (Histone-2B CFP) and a marker for actin (Actin Chromobody tagRFP), allowing us to image the orientation of cell divisions relative to apical surfaces, marked with strong actin staining. We imaged cysts grown on Matrigel using the Opera Phenix High Content imaging microscope, imaging at 7.5 min intervals over periods of 6-12 hours. In control cysts (see top panels in Figure 2k, Supplementary Movie 1) cells tended to divide in the plane of the epithelium, when measured at either metaphase (Figure 2l) or anaphase to early cytokinesis (Supplementary Figure 2l). In cysts with constitutive knockdown of either CASK or Dlg1, we saw a randomisation of the angle of cell division at both metaphase and anaphase/early cytokinesis (Figure 2k, quantification in Figure 2l, Supplementary Figure 2l; see also Supplementary Movies 2 and 3). Together with the results from fixed imaging, this indicates that both CASK and Dlg1 are required for normal epithelial spindle orientation.

Further, we used our live imaging data to determine the time required for cells to progress from prophase through to early cytokinesis, and found that there was no change in mitotic progression in CASK and Dlg1 knockdown cysts compared to control cysts (Supplementary Figure 2m). This indicates that there is no spindle orientation checkpoint in MDCKII cells.

### The direct interaction between CASK and Dlg1 is required for correct spindle orientation

Having seen similar results with respect to lumen formation and spindle orientation upon depletion of either CASK or Dlg1, we set out to investigate further the relationship between these two proteins, and determine whether CASK affects spindle orientation directly through its interaction with Dlg1. The most N-terminal amino acids of Dlg1 have been identified as those required for binding to CASK [14]; this region interacts with the more N-terminal of two L27 domains in CASK (Figure 3a) [14]. We reasoned that over-expression of this domain might be sufficient to saturate the Dlg1-binding site on CASK, and thereby block the endogenous CASK-Dlg1 interaction. We generated a doxycycline-inducible plasmid containing the first 66 amino acids of Dlg1 fused to an HA tag (Figure 3a) and made stable MDCKII cells containing this construct, which we named D66- HA. Similarly to other small interfering constructs we have previously generated [22] we were not able to detect the D66-HA peptide directly by western blotting. However, upon treatment with doxycycline HA expression could be detected at the cell membrane (Figure 3b), indicating that the construct is expressed and localises similarly to endogenous Dlg1. Next, we tested the capacity of this construct to interact with CASK. We performed an immunoprecipitation in the presence or absence of doxycycline and saw that only in the doxycycline treated cells was the HA antibody able to co-immunoprecipiate endogenous CASK (Figure 3c). This indicates that the D66-HA construct is sufficient to interact with CASK in cells. To determine whether expression of the D66-HA construct is able to disrupt the normal CASK-Dlg1 interaction, we immunoprecipitated endogenous CASK and probed for Dlg1. In cells treated with doxycycline there is a marked decrease in the amount of Dlg1 immunoprecipitated (Figure 3d), indicating that expression of the D66-HA peptide is able to block the normal CASK-Dlg1 interaction. To test whether interaction with CASK was required for Dlg1 membrane localisation, we stained MDCKII cysts for Dlg1 and saw that expression of the D66-HA peptide caused a loss of Dlg1 staining at the basolateral membrane in these cysts (Figure 3e).

**Figure 3-.**
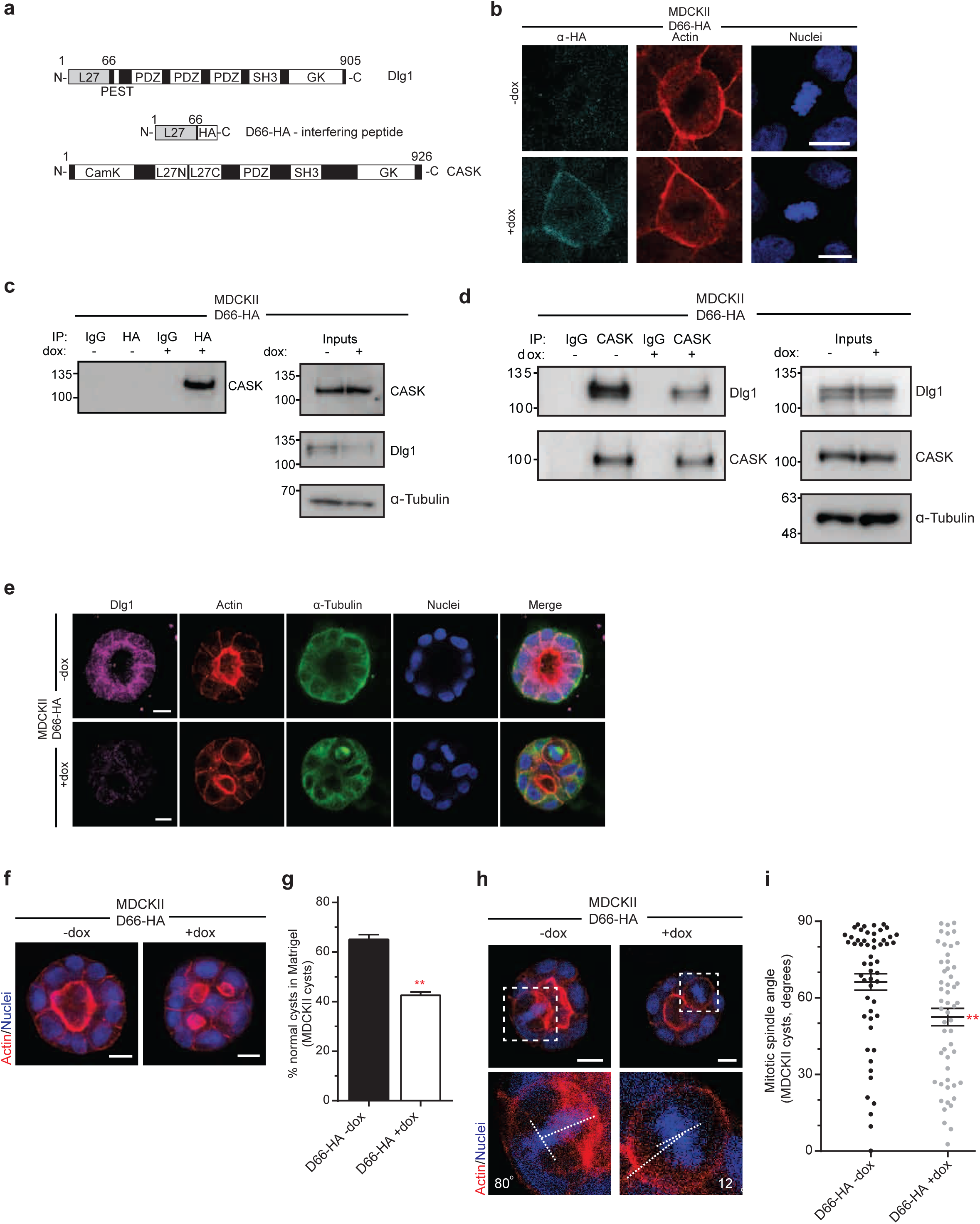
The interaction between CASK and Dlg1 regulates spindle orientation and lumen formation. a) Schematic of the protein structure of Dlg1 and CASK, showing in grey the N-terminal domain of Dlg1 which binds to the L27N domain of CASK. The interfering peptide, D66-HA, is composed of the N-terminal domain of Dlg1 followed by an HA tag. b) Widefield images showing localisation of D66- HA at the cell membrane of metaphase cells, when D66-HA is expressed following dox treatment (left panels). c) Western blot for CASK following immunoprecipitation by HA from lysates of MDCKII cells inducibly expressing the D66-HA interfering peptide following addition of doxycycline (+ dox). d) Endogenous CASK was immunoprecipitated from MDCKII cell lysates and co-precipitated endogenous Dlg1 was detected by immunoblotting. Dlg1 pull-down is reduced following expression of D66-HA (+dox) for 1 day. A representative western blot from three independent experiments is shown. e) Images of MDCKII cysts grown in Matrigel showing reduction of basolateral Dlg1 staining following expression of D66-HA. f) Representative images of cysts grown in Matrigel from MDCKII cells inducibly expressing the D66-HA interfering peptide; D66-HA was expressed with the addition of doxycycline (+dox), leading to a multilumen phenotype. g) Quantification of normal cysts following D66-HA expression (+dox). N=8/9 independent experiments, >100 cysts quantified per experiment; ** p= 0.002 (Mann-Whitney Test). Values are mean +/- SEM. h) Representative images of metaphase cells of MDCKII cysts with (+dox) or without (-dox) D66-HA expression. Insets annotated to show angle of cell division relative to the apical surface. i) Quantification of spindle orientation with and without the expression of D66-HA. n=54/53 from three independent experiments; ** p=0.004 (Mann-Whitney Test). Error bars show mean +/- SEM.

Next we examined the physiological effects of blocking the CASK-Dlg1 interaction. We grew MDCKII cells containing the D66-HA peptide in 3D culture, and saw that doxycycline treatment increased the number of multilumen cysts compared with untreated cells (Figure 3f and 3g). To test whether the CASK-Dlg1 interaction is required for normal spindle orientation, we measured the angle of cell divisions in 3D in cells expressing D66-HA, and saw a randomisation of the spindle angle compared with control cysts (Figure 3h and 3i). Expression of this construct was also able to disrupt the orientation of the earliest cell divisions when cells were grown in collagen (Supplementary Figure 3a, 3b), indicating that the CASK-Dlg1 interaction is important even in a context where cells lack lateral neighbours. Together these results indicate that the direct interaction between CASK and Dlg1 is required for the localisation of Dlg1 at the basolateral membrane, and for normal growth of epithelial structures in 3D.

### The CASK-Dlg1 interaction is required for localisation of the canonical LGN-NuMA complex

Spindle orientation requires a conserved set of proteins including LGN, which is normally localised to the basolateral membrane in epithelial cells. LGN localisation to the membrane is mediated in part by Gαi binding, but additional factors are also required for correct LGN localisation and function [23]. We therefore tested whether LGN localisation was affected by depletion of CASK. We stained 2D cultures of MDCKII cells and saw clear membrane localisation of LGN in control metaphase cells, as well as localisation to the spindle poles (Figure 4a). Upon CASK knockdown, membrane staining of LGN was lost, whereas the spindle pole localisation was unaffected (Figure 4a). We quantified this effect by measuring the ratio of membrane to cytoplasmic LGN (Supplementary Figure 4a), and saw a significant decrease in LGN membrane localisation in CASK depleted cells (Figure 4b). Dlg1 has been implicated in LGN localisation in neuroepithelial cells; we saw a significant decrease in the membrane to cytoplasmic ratio of LGN in Dlg1 depleted cells (Figure 4a and 4b). In some cells, however, LGN localisation depends on laterally-localised E-Cadherin [24], and E-Cadherin is required for correct spindle orientation in mouse prostate epithelial cells [25]. To test whether Dlg1 acts through localisation of E-Cadherin, we stained MDCKII cells for E-Cadherin following both CASK and Dlg1 knockdown. We did not see a difference in E-Cadherin staining following loss of CASK or Dlg1 (Supplementary Figure 4b). This suggests that depletion of Dlg1 or CASK do not disrupt spindle orientation through regulation of E-Cadherin localisation or by substantive changes to adherens junctions, but instead by a process dependent on direct interaction between Dlg1 and LGN.

**Figure 4-.**
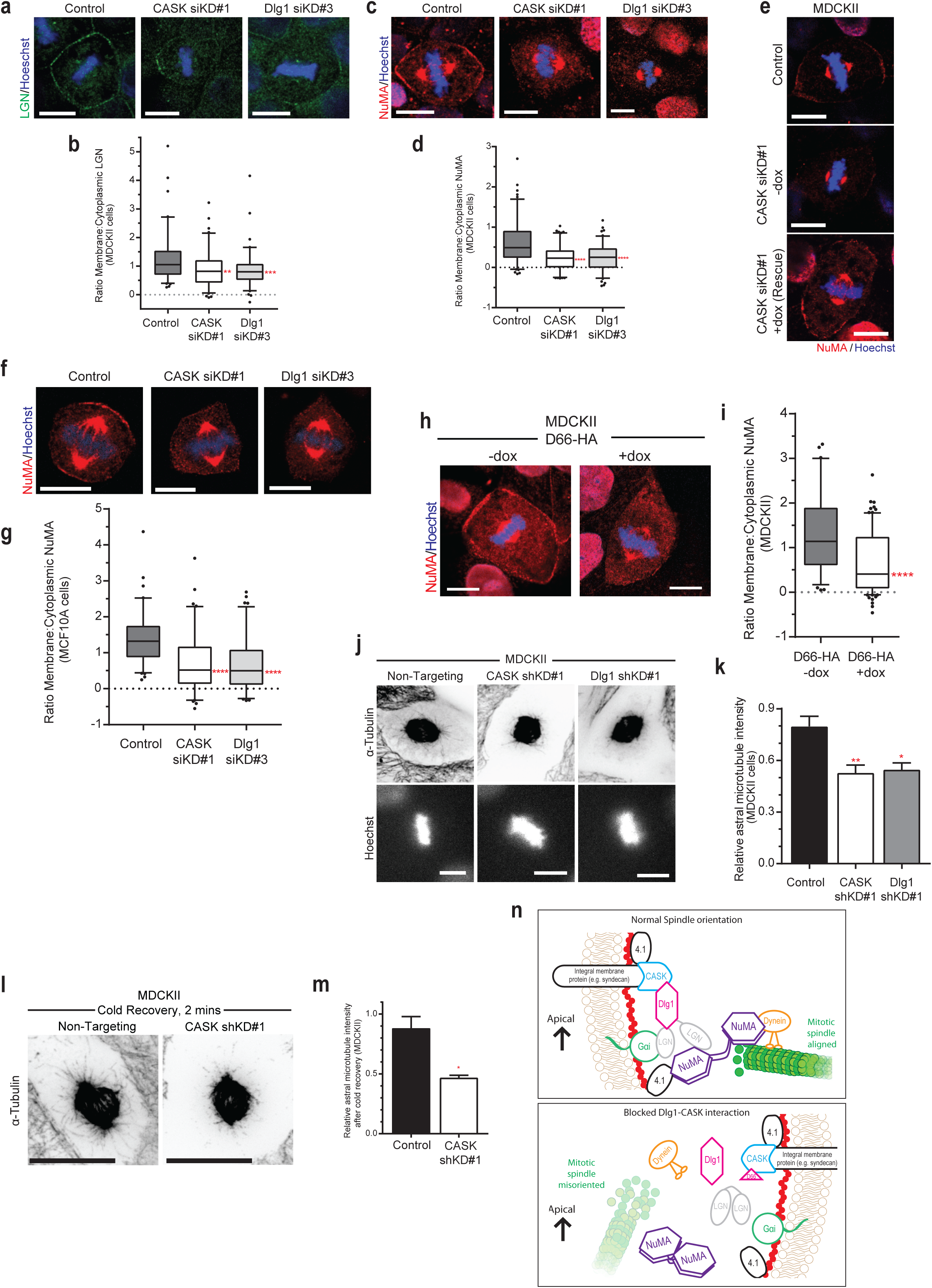
CASK and Dlg1 localise the spindle orientation complex of LGN and NuMA. a) Representative confocal images of LGN staining in Control cells (Non-Targeting siRNA), with reduced membrane-associated LGN in CASK and Dlg1 knockdown cells (CASK siKD#1 and Dlg1 siKD#3). b) Quantification of Membrane:Cytoplasmic LGN ratio. n=86/87/90 membrane measurements from three independent experiments; ** p=0.0013, *** p=0.0005 (one-way Anova). Box shows 25-75 percentile marked with the median, whiskers show 5-95 percentile, dots represent measurements outside this range. c) Representative confocal images of NuMA staining in Control cells (Non-Targeting siRNA), with reduced membrane-associated NuMA in CASK and Dlg1 knockdown cells (CASK siKD#1 and Dlg1 siKD#3). d) Quantification of Membrane:Cytoplasmic NuMA ratio. n=106/108/110 membrane measurements from three independent experiments; **** p=4.9×10^−11^ and 2.2×10^−10^(one-way Anova) for CASK siKD#1 and Dlg1 siKD#3 respectively. Box shows 25-75 percentile marked with the median, whiskers show 5-95 percentile, dots represent measurements outside this range. e) Representative confocal images of NuMA staining in MDCKII cells inducibly expressing siRNA-resistant CASK upon addition of dox (+dox) treated with either Non-Targeting (Control) or CASK siKD#1. f) Representative confocal images of NuMA staining in Control MCF10A cells (Non-Targeting siRNA), with reduced membrane-associated NuMA in CASK and Dlg1 knockdown MCF10A cells (CASK siKD#1 and Dlg1 siKD#3). g) Quantification of Membrane:Cytoplasmic NuMA ratio. n=84/81/88 membrane measurements from three independent experiments; **** p=3.7×10^−7^and 9.0×10^−9^ (one-way Anova) for CASK siKD#1 and Dlg1 siKD#3 respectively. Box shows 25-75 percentile marked with the median, whiskers show 5-95 percentile, dots represent measurements outside this range. h) Confocal images of MDCKII cells in metaphase stained for NuMA, showing a decrease in membrane localised NuMA upon expression of D66-HA. i) Quantification of Membrane:Cytoplasmic NuMA ratio in D66-HA expressing (+dox) and non-expressing (-dox) cells, n=77/91 membrane measurements from three independent experiments; **** p=1.4×10^−6^ (unpaired Student’s t-test, two tailed). Box shows 25-75 percentile marked with the median, whiskers show 5-95 percentile, dots represent measurements outside this range. j) Spinning disc confocal maximum intensity projections of metaphase cells stained with α- Tubulin showing long astral microtubules in control cells (Non-Targeting shRNA), which are reduced or absent in CASK shKD#1 and Dlg1 shKD#1 knockdown cells. Images are inverted to more clearly display the astral microtubules. k) Quantification of astral microtubule intensity (N=8/7/4 independent experiments, >20 cells quantified per experiment). Values are mean +/- SEM. ** p=0.004, * p= 0.0217. All scale bars are 10 µm. l) Representative images of MDCKII cells following two minutes recovery from cold treatment. Maximum intensity projections of spinning disc confocal stacks. m) Quantification of astral microtubule intensities from cold recovery experiments. N=3; * p= 0.0144 (unpaired Student’s t-test, 2 tailed). Values are mean +/- SEM. Data were analysed using an unpaired Student’s t-test, two tailed. n) Proposed model for the role of the CASK-Dlg1 interaction in localising LGN and NuMA to the lateral membrane and providing a positional cue for the correct orientation of the mitotic spindle.

LGN binds to NuMA, and helps localise it to the cell membrane [26, 27]. We therefore tested whether CASK and Dlg1 are required for normal NuMA localisation. We stained our 2D MDCKII cultures with a NuMA-specific antibody and in control metaphase cells we saw clear staining at the membrane, as well as at the mitotic spindle (Figure 4c). In both CASK and Dlg1-depleted cells, we saw a striking reduction in membrane staining of NuMA, while the spindle still stained strongly (Figure 4c). We quantified the membrane to cytoplasmic ratio of NuMA and saw a significant decrease following depletion of either CASK or Dlg1 (Figure 4d). We were able to restore membrane localisation of NuMA in our CASK rescue cells (Figure 4e), demonstrating that this effect is specific to loss of CASK. We observed a similar decrease in the membrane to cytoplasmic ratio of NuMA following siRNA knockdown of CASK and Dlg1 in MCF10A cells in 2D culture (Figure 4f and 4g), demonstrating a conserved role for both CASK and Dlg1 in NuMA localisation. To test whether the CASK-Dlg1 interaction itself is required for the correct localisation of NuMA, we stained cells expressing the interfering D66-HA peptide, and saw a loss of membrane NuMA localisation and a significant reduction in the membrane to cytoplasmic ratio of NuMA (Figure 4h and 4i). This indicates that the direct interaction between CASK and Dlg1 is necessary for normal membrane localisation of NuMA, a key component of the spindle orientation machinery.

NuMA is able to interact with microtubules which connect the mitotic spindle to the plasma membrane [28] and provide a pulling force to align the mitotic spindle [1]. We therefore wanted to see whether there was an effect on astral microtubules in cells depleted for CASK or Dlg1. We stained 2D MDCKII cultures for alpha-Tubulin, and imaged metaphase cells to look at astral microtubules. We observed a reduction in astral microtubules in CASK or Dlg1 depleted cells; we quantified the relative astral microtubule intensity relative to microtubule staining at the mitotic spindle (Supplementary Figure 4c) and saw a significant reduction in both CASK and Dlg1 depleted cells compared with control cells (Figure 4j and 4k). To test whether this was due to a general reduction in astral microtubule numbers, or to an effect on microtubule stability, we subjected cells to cold treatment, which depolymerises microtubules, and then returned them to 37°C to allow microtubule regrowth. A significantly higher regrowth was seen in control cells compared to CASK depleted cells, indicating a defect in microtubule dynamics following CASK knockdown (Figure 4l and 4m). We wondered whether this defect in microtubule dynamics would mean that there would be less movement of the mitotic spindle in CASK and Dlg1-depleted cells. We returned to our live imaging data (Figure 2k) and for those individual cells where we were able to measure the angle of the mitotic spindle at metaphase relative to the apical surface of the cell, and then again later (as in the example images in Figure 2k), we measured the amount of spindle rotation between metaphase and anaphase/early cytokinesis. Movement that increased alignment was recorded as positive, movement which decreased the alignment was recorded as negative. As shown in Supplementary Figure 4d, most of the movement is positive, indicating on-going processes to align the cell division plane after metaphase, even in cells depleted of CASK and Dlg1. There is a similar amount of movement in control and CASK or Dlg1 depleted cells, suggesting that the defect in spindle orientation is not due to an overall lack of spindle movement. We conclude that CASK and Dlg1 affect the alignment of the mitotic spindle during or prior to metaphase, rather than the absolute ability of the mitotic spindle to rotate.

Correct spindle positioning is vital for oriented cell division in epithelia [9]. We have identified a novel role for the MAGUK protein CASK in spindle orientation in epithelial cells, and we propose that CASK acts as an upstream factor for Dlg1 localisation. Downstream of CASK and Dlg1 are the conserved proteins of the spindle orientation complex, LGN and NuMA, whose localisation depends on membrane-bound Dlg1 (see model in Figure 4o). CASK contains a hook domain which allows for interaction with the membrane via 4.1 proteins as well as by binding to Syndecans [29], and therefore provides the link for localisation of Dlg1, which does not itself bind membranes. Loss of CASK phenocopies the effects of directly depleting Dlg1, showing the close relationship between these two proteins. Moreover, when we block the binding using an interfering peptide we recapitulate the effects of depleting either CASK or Dlg1, indicating that direct interaction is required for Dlg1 localisation and function.

Dlg1 has been linked to NuMA localisation at tricellular junctions in fly epithelia, [30], as well as to both LGN and NuMA localisation in HeLa cells [6]. However, it is apparent that oriented cell divisions are regulated differently in different cell types in different organisms [1, 31]. Here we show that this role for Dlg1 is conserved in untransformed mammalian epithelia. Importantly we show that both CASK and Dlg1 are required for spindle orientation and lumen formation. This requirement for Dlg1 may explain in part the multilumen phenotype seen in SHIP2-depleted MDCKII cells which also mislocalise Dlg1 [32], as well as in prostate epithelial depleted of E-Cadherin where Dlg1 is lost from the membrane due to disruption of adherens junctions [25].

Mutations in CASK cause mental retardation as a result of microcephaly [18]. Microcephaly is a common consequence of misoriented cell divisions in neuronal progenitor cells due to an imbalance in the production of differentiated neurons and self-renewal of progenitors [33] and it would be interesting to determine whether the role for CASK in spindle orientation is conserved in neuronal stem cells. In neurons, CASK binding to Dlg1 is able to alter the conformation of Dlg1, which in turn changes the binding targets of Dlg1 [20]. While the loss of Dlg1 from the membrane following CASK knockdown may be sufficient to explain the misorientation of the mitotic spindle, it is interesting to speculate that CASK binding may also mediate the conformation of Dlg1 to facilitate LGN binding.

We see a reduction in astral microtubule density following CASK or Dlg1 knockdown, which we hypothesise is due to changes in the rate of capture at the membrane following loss of NuMA localisation. Other studies have reported that changes in astral microtubule number and growth are able to affect spindle orientation; for instance CYLD stabilises microtubules and promotes assembly of the spindle orientation machinery at the membrane [34]. Loss of this CASK-Dlg1 complex leading to changes in astral microtubule dynamics suggests the presence of a feedback mechanism which may merit further investigation.

Loss of spindle orientation and the genesis of multilumen structures have been linked on a number of occasions, and we see that our multilumen structures retain polarity at the cellular level while having lost the overall polarity of the cyst. This is reminiscent of the early stages of many tumours, including DCIS [35], where aberrant structures are contained within a single basement membrane. Interestingly low levels of CASK mRNA are prognostic of poor outcomes in breast cancer (kmplotter.org) and disruption of the CASK-Dlg1 interaction (through loss of the SH3 domain of Dlg1) [36] has been reported to impair normal kidney development and lead to the formation of cystic kidneys [36]. Studies in fruit flies indicate that loss of spindle orientation can be compensated for by reintegration of the misoriented cells back into the epithelium [37]. Through our live imaging results we saw a range of outcomes for individual misoriented divisions, including reintegration of daughter cells in the epithelium, as well as formation of new luminal structures (Supplementary Movies 2, 3). However the resolution of imaging in three dimensions was insufficient to build a quantitative picture of the outcome of these events. Nevertheless, in line with previous reports [23], we can detect no spindle orientation checkpoint as oriented and misoriented divisions proceed at the same rate in our live imaging experiments. This inability for cells to correct misoriented divisions highlights the importance of correctly establishing polarity prior to entry into mitosis.

## Methods

### Antibodies

Working dilutions of antibodies for immunoblotting (IB) and immunofluorescence (IF) are shown below.

α-Tubulin [DM1A] – Abcam ab7291 Mouse (1:2500 IF, Methanol, 1:5000 WB); β-actin – Sigma Clone AC-15, Mouse (1:10,000 WB); E-cadherin – Abcam DECMA-1 Rat (1:200 IF, Methanol); GP135/Podocalyxin (IF: 1:2000, Formaldehyde) – gift from Dave Bryant, Beatson Institute for Cancer Research; Actin – Alexa Fluor^®^ 488/568 Phalloidin - Molecular Probes; CASK – CASK (C-6) sc-13158 mouse (1:100 IF, 1:1000 WB, 3 µg/IP); Dlg1 (SAP97) (H-60): Santa Cruz, sc-25661 Rabbit (1:100 IF, Methanol; 1:1000 WB); β-catenin – Upstate, 06-734, 1:200 (IF); Anti-NuMA antibody (ab36999) Abcam (IF: 1:150, Methanol); LGN (Millipore, ABT-174, (IF: 1:250, Methanol); Pericentrin (PRB-432C) Covance (IF: 1:2000, Methanol).

Secondary antibodies: IgG-peroxidase-conjugated (IB 1:5,000, GE Healthcare); EasyBlot anti Mouse IgG (HRP), GeneTex (for reprobing IP membranes, also using Easy Blocker solution (GeneTex); Alexa Fluor 488, 568, 647-conjugated (IF 1:500, Molecular Probes).

### Cell culture

All cells were cultured in a 37 °C, 5% CO2 incubator. Parental MDCKII (from ECACC, operated by Public Health England) were maintained in Dulbecco’s Modified Eagle Medium (DMEM, Invitrogen) in the presence of 10% fetal bovine serum (FBS, GIBCO). Cell lines were routinely tested for mycoplasma contamination by our in-house facility. MDCKII with dox-inducible Myc-CASK were maintained in DMEM with 10% tetracycline-free FBS with G418 and puromycin (2 μg ml-1, Sigma). MDCKII cells expressing histone-2B-GFP and Actin Chromobody were grown in DMEM with 10% tetracycline-free FBS with G418 and blasticidin (5 μg ml-1, Sigma). MCF10A cells (a gift from Dr Gillian Farnie, University of Oxford), were cultured in complete medium composed of the whole minimal DMEM medium components supplemented with 100 ng/ml cholera toxin, 20 ng/ml epidermal growth factor, 10 μg/ml insulin, and 0.5 μg/ml hydrocortisone.

### Cysts

Two methods for 3D culture were used in this paper. A) Base layers (250 μl) of Collagen I solution (2mg/ml Collagen I with added 1M NaOH at 0.0023 times the volume of added Collagen I) were set in 24-well plates (20 min, 37 °C), before the addition of 300 μl collagen top-layers containing 3.6×10^4^ cells for lumen formation or spindle orientation assays or 2×10^5^ cells for early spindle orientation assays. Gels were cultured in 700 μl medium for 1 day for early spindle orientation assays, 4-6 days for spindle orientation assays, and 10 days for lumen formation assays before immunostaining, with 50% of the media being replaced every 2-3 days.

Cyst-containing gels were washed twice in PBS; fixed in 3.7% formaldehyde for 15 mins at room temperature; washed twice in PBS; permeabilised in 0.5% Triton/PBS for 30 min, room temperature; washed twice in PBS; incubated with blocking solution (10% FBS/PBS) for 1 h, room temperature; incubated with primary antibody in 2% FBS/PBS overnight, 4 °C; washed in PBS 6×30 min, 4 °C; incubated with secondary antibody, Phalloidin and Hoechst 1% FBS/PBS overnight, 4 °C; washed in PBS 6×30 min, 4 °C; and then mounted onto slides in mounting medium for imaging on the low-light, macro-confocal or SP8 confocal microscopes.

B) Cells were trypsinised to a single cell suspension at 1.5 × 10^4^ cells/ml in complete medium containing 2% Matrigel (BD Matrigel Matrix Phenol Red-Free 10 ml 356237). Suspensions (250 μl) were plated into 8-well coverglass chambers (Nunc), pre-coated with 10 μl of 100% Matrigel. Cells were grown for 4 days before fixation in 3.7% formaldehyde for 15 mins at room temperature; permeabilised in 0.5% Triton/PBS for 20 min, washed twice in PBS; incubated with blocking solution (10% FBS/PBS) for 1 h, and stained as above. A similar method was applied for the formation of MCF10A cysts, using the same reagents and cell plating density. These cysts were left to grow for 10 days in Matrigel culture. 50% of media was replaced every 2-3 days.

### Generation of cell lines

Plasmids were introduced into cells either by transfection using TransIT-LT1 (Mirus) according to the manufacturer’s instructions or by retroviral transduction as previously described [38]. For inducible overexpression, MDCKII were retrovirally transduced with pRetro-Tet-ON followed by selection with G418 (1 mg ml^−1^, Sigma). pRetro-XT-based constructs were then retrovirally transduced and cells selected with puromycin (2 μg ml^−1^, Sigma Sigma). For live imaging, cells were FACS sorted to produce a population expressing both CPF and RFP markers. For inducible cell lines, clones of MDCKII cells expressing dox-inducible shRNAs to Cask, against two independent target sequences (Cask RNAi#1 and #2), were selected by single cell sorting and screening for efficient knockdown of CASK upon addition of Doxycycline.

### Plasmids and cloning

#### CASK WT

The EcoRI myc-Cask fragment from myc-Cask-FL (a gift from Yi-Ping Hsueh) was inserted into the pRetro-X-tight(puro) (Clontech), and mutated by Quikchange II mutatgenesis (Stratagene) to generate a sequence resistant to CASK shKD#1 shRNA and CASK siKD#1 siRNA sequences. The changes were as follows:

Wild-type sequence: TTA AGT ACA GAA GAT C

Leu Ser Thr Glu Asp

Resistant sequence: TTG AGC ACC GGG GAC C

The following primers were used for the mutagenesis:

Forward: 5’- CATCAAGTCCAGGGTTGAGCACCGGGGACCTAAAGCGGGAAGC – 3’

Reverse: 5’ – GCTTCCCGCTTTAGGTCCCCGGTGCTCAACCCTGGACTTGATG – 3’

#### D66HA

The N-terminal region of Dlg1 was amplified by PCR, and a HA tag added to the sequence of the reverse primer, along with restriction sites for insertion into the pRetro-X-Tight(puro) vector (Clontech). Correct insertion was verified by Sanger sequencing. Primers were:

Dlg66 Forward: 5’-ACGTGCGGCCGCCTTCCTGATTCTGGA-3’

Dlg66_Reverse: 5’- ACGTCTTAAGTCAAGCGTAATCTGGAACATCGTATGGGTATGGTTCACACTGCTTTGAATGA-3’shRNAs were screened for their Cask knockdown ability by transient transfection in HEK293T cells and two sequences that successfully down-regulated Cask (CASK iKD#1 and CASK iKD#2) were further sub-cloned into the pA’-TO dox-inducible RNAi vector (as previously described in [39]). Additionally, the shRNA-targeting sequence from CASK iKD#1 was cloned into the pRetro-Super (Clontech) vector, for constitutive knockdown of CASK. For Dlg1 knockdown, we used the sequence si1 against Dlg1 (from [32]) to produce primers to clone into pRetro-Super to generate Dlg1 shKD#1.

Primers were as follows:

CASK iKD#1 forward:

5’-GATCCCGTTAAGTACAGAAGATCTATTCAAGAGATAGATCTTCTGTACTTAACTTTTTGGAAA-3’

CASK iKD#1 reverse:

5’-AGCTTTTCCAAAAAGTTAAGTACAGAAGATCTATCTCTTGAATAGATCTTCTGTACTTAACGG-3’

CASK iKD#2 forward:

5’-GATCCCGCTCAGATGGAATGCTTTATTCAAGAGATAAAGCATTCCATCTGAGCTTTTTGGAAA-3’

CASK iKD#2 reverse:

5’-AGCTTTTCCAAAAAGCTCAGATGGAATGCTTTATCTCTTGAATAAAGCATTCCATCTGAGCGG-3’

CASK shKD#1 forward:

5’-GATCCGATAAGTACAGAACATCTATTTCAAGAGAATTCTTCT-3’

CASK shKD#1 reverse:

5’-AATTCAAAAAATTAAGTACAGAAGATCTATTCTCTTGAAATAGAT-3’

Dlg1 shKD#1 forward:

5’-GATCCCGATATCCTCCATGTTATTATTCAAGAGATAATAACATGG-3’

Dlg1 shKD#1 reverse:

5’-AGCTTTTCCAAAAAGATATCCTCCATGTAATAATCTCTTGAATAA-3’

The Non-Targeting (shRNA knockdown) plasmid used targets the sequence: 5’-ATGAAGTCGCATGGTGCAG-3’

The Actin Chromobody was purchased from ChromoTek. The Histone-2B-CFP plasmid was the same as previously used in [39].

### Transient transfection of siRNA

Transient silencing of CASK and Dlg1 was achieved by transfection of siRNA oligos from Eurofins MWG operon into MDCKII cells using Lipofectamine RNAiMax (Invitrogen, Life Technologies) according to the manufacturer’s instructions. Cells were processed and analysed 48 h post transfection. siRNA sequences were as follows:

CASK siKD#1 5’-GGUUAAGUACAGAAGAUGUAATT-3’

Dlg1 siKD#1 5’-CAACTCTTCTTCTCAGCCTTT-3’

Dlg1 siKD#2 5’-CAGAAGAACAAGCCAGAAATT-3’

sDlg1 siKD#3 5’-GAUAUCCUCCAUGUUAUUATT-3’

Non-Targeting: Dharmacon—Non-Targeting siRNA #4

### Protein analysis

Cells were lysed in IP lysis buffer (50 mM Tris-HCl, pH 7.5, 150 mM NaCl, 1% (v/v) Triton-X-100, 10% (v/v) glycerol, 2 mM EDTA, 25 mM NaF and 2 mM NaH2PO4) containing a protease inhibitor cocktail (Sigma) and phosphatase inhibitor cocktails 1 and 2 (Sigma).

For immunoprecipitation, lysates were incubated with the appropriate antibody pre-bound to 20 μl of GammaBind G Sepharose (Amersham) for 3 h at 4 °C (for HA pull-down) or overnight (for endogenous IPs), washed in IP lysis buffer, then eluted with 1 × SDS–PAGE sample buffer (Nupage, Invitrogen).

### Immunofluorescence – 2D culture

For immunofluorescence in 2D, cells were grown on coverslips and fixed with either 100% ice-cold methanol for 5 min at −20 °C or 3.7% formaldehyde for 20 min at room temperature. Staining was performed by permeabilisation in 0.5% triton in PBS and blocking in 1% BSA in PBS before successive incubation with primary and then secondary antibodies. Coverslips were mounted to slides using Fluromount-G (Southern Biotech) along with Hoescht (1:5000) for nuclear staining.

### Immunofluorescence – 3D culture

For MDCKII cysts grown in collagen, the collagen plug was fixed in-well with Formaldehyde for 20 minutes at room temperature, and washed three times with PBS. Cysts were permeabilised with 0.5% triton in PBS for 20 minutes at room temperature with gentle rocking, before the whole collagen plug was gently placed in an Eppendorf, before being blocked with 2% BSA for one hour. Cysts were left in 1% BSA in PBS containing primary antibodies overnight at 4 degrees with gentle rotation, before being washed 6 times for 30 minutes each time. Secondary antibodies were applied overnight in the same way. The collagen plug was then mounted on a glass slide, covered with mounting media (Fluromount-G, Southern Biotech) and a coverslip gently placed on top.

For MDCKII cysts and MCF10A cysts grown in Matrigel, these were fixed in the wells of the chamber slide using formaldehyde, as above, stained in the same way as for collagen-grown cysts.

### Microscopy

The spinning-disc confocal microscope was based around an Olympus IX81, using the Sedat filter set (Chroma, 89000), an array of imaging lasers (406, 488, 548, 645 nm) and an Apo ×100 1.45 NA oil objective. The low-light microscope was based around a Zeiss Axiovert 200M, with an Andor iXon DU888+ camera, a 300 W xenon light source, using the Sedat filter set, and a Zeiss alpha plan ×100 1.45 NA oil objective. The macro-confocal microscope was based around a Leica TCS LSI using a Leica plan Apo ×5 0.50 NA objective, an array of solid-state imaging lasers (488, 532 and 635 nm) and an ultrahigh dynamic photomultiplier. Images were captured using Metamorph software (Molecular Devices) or Leica proprietary imaging software. The Deltavision Core (Applied Precision Instruments) system was based around an Olympus IX71 microscope with illumination achieved by white light LED and a 300 W Xenon light source for fluorescence. The Sedat filter (Chroma, 89000) set was utilized for fluorescence imaging using a UPLSAPO 60XO 1.35NA objective, and image capture was via a Roper Cascade II 512B EMCCD camera and SoftWorx software (Applied Precision Instruments). Images were also acquired on an inverted Leica TCS SP8 confocal microscope equipped with PMT and Hybrid (HyD) detectors, with the tunable white laser (WLL) for Alexa Fluor 488, Alexa Fluor 555 and Alexa Fluor 647, and 405 nm UV laser for Hoescht. For 2D imaging, the x63 1.45 NA oil immersion objective was used, while for 3D imaging the 25x xx NA water lens was used. For 3D imaging we also used an upright Leica TCS SP8 confocal microscope equipped with PMT and Hybrid (HyD) detectors, and laser lines at 405, 488 568 and647. Images were captured in LAS AF (3.0.1) Leica software.

### Live imaging 3D culture

Live imaging was performed using cells in 96-well plates and imaged using an Opera Phenix (Perkin Elmer) with temperature and environmental controls (37 °C and 5% CO2), using a × 20 NA 0.95 water immersion lens (Zeiss). Frames were captured at 7.5 min intervals for between 6 and 12 hours. Movies were exported from the Columbus software and spindle orientation angles were analysed manually in ImageJ.

### Quantification of LGN and NuMA levels

From confocal stacks of mitotic cells, the fluorescence of LGN and NuMA at the membrane was calculated for each pole of the cell, in the stack where the fluorescence at the membrane was strongest, by manually drawing a region of interest in ImageJ along the membrane (line width:5), and recording the mean intensity. A second line was drawn, adjacent to the first, and of similar length, in the cytoplasm immediately adjacent to the membrane, and the intensity recorded. The ratio of Membrane:Cytoplasm intensity was then calculated in Microsoft Excel and imported into Prism for analysis.

### Quantification of astral microtubule intensity

Maximal projections of spinning disc confocal stacks were generated using ImageJ. Two regions of interest were drawn, encompassing a) the mitotic spindle and b) the cell area containing astral microtubules. The formula (b-a)/a was used to calculate the relative astral microtubule intensity, normalised to the intensity of the mitotic spindle.

### Quantification of spindle orientation

For spindle orientation in 3D culture, confocal or Widefield images of the dividing cell were manually analysed using the Measure Angle tool in ImageJ, with a line drawn through the axis of the dividing cell, and the angle measured to the nearest apical surface. For 2D spindle orientation, the xyz coordinates of the centre of each centrosome (marked by Pericentrin staining) were determined by marking the appropriate stacks when viewed in ImageJ. Standard trigonometric functions were used to calculate the angle relative to the surface of the microscope slide.

### Statistical analysis

Statistical differences between data were analysed in Prism (GraphPad software) using Anova or two-tailed unpaired student’s t-test as appropriate (i.e. comparison of only two groups) (P values are specified in figure legends).

## Supporting information

## Acknowledgements

We thank Gillian Farnie for her help in establishing the MCF10A 3D culture system. We thank Dave Bryant for the kind gift of the GP135/Podocalyxin antibody, and Yi-Ping Hsueh for the kind gift of the full-length, myc-tagged CASK plasmid. We thank Adam Hurlstone and members of the Cell Signalling lab for their critical reading of the manuscript. We thank the Molecular Biology Core Facilities and the Advanced Imaging Group at the Cancer Research UK Manchester Institute for their assistance with sequencing and microscopy. This work was supported by Cancer Research UK (grant number C5759/A12328) and Worldwide Cancer Research (grant number 12-0037).

## Supplementary Figure Legends

**Supplementary Figure 1.**
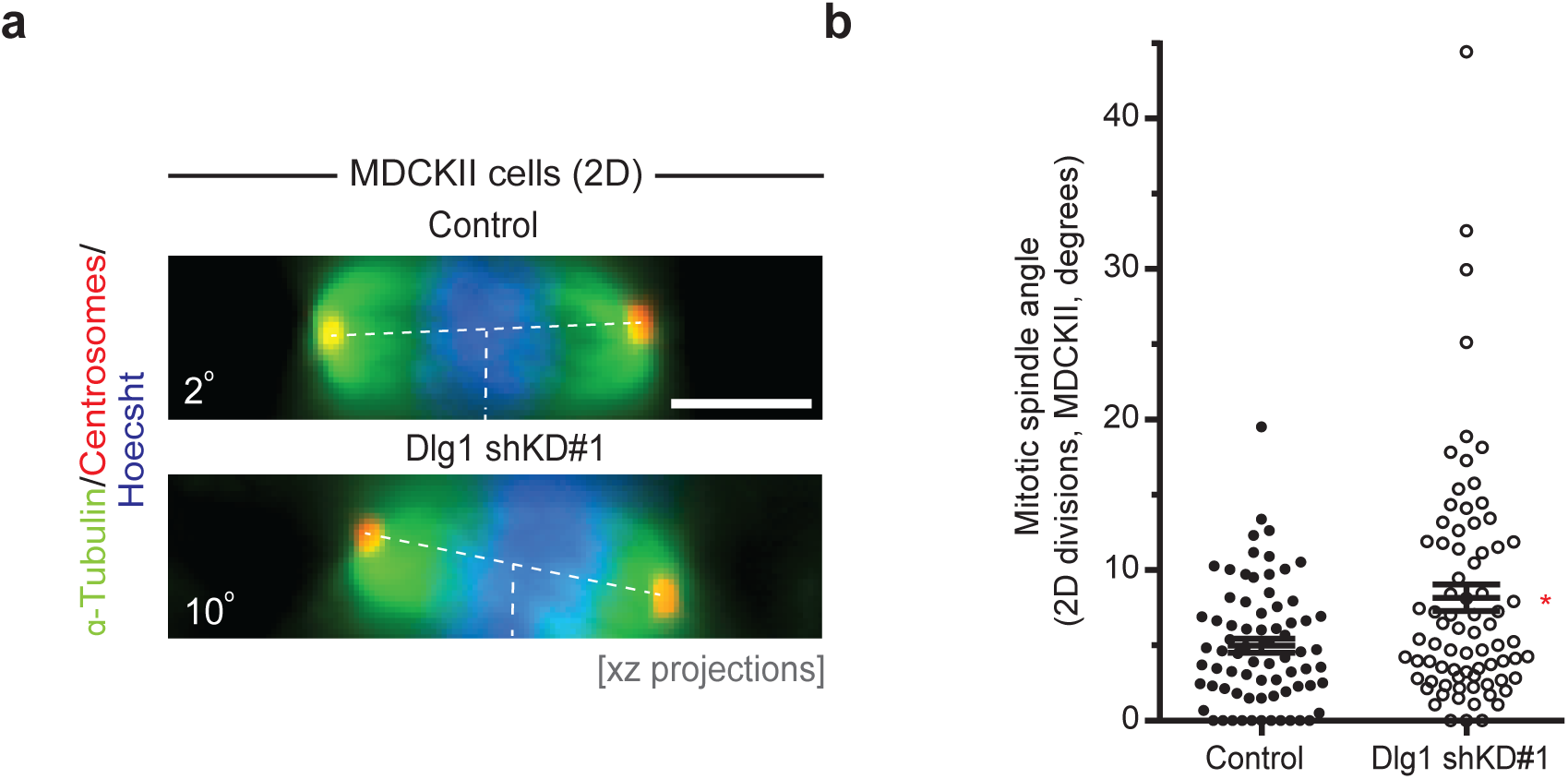
Representative images of XZ projections from MDCKII cells grown in 2D, showing tilted mitotic spindles at metaphase after Dlg1 knockdown, and annotated to show guide lines and spindle angles. Quantification of spindle angles from MDCKII Control (Non-Targeting) and Dlg1 knockdown cells grown in 2D, n=73/77 from three independent experiments; * p= 0.033 (Kruskal-Wallis test). Error bars show mean +/- SEM.

**Supplementary Figure 2.**
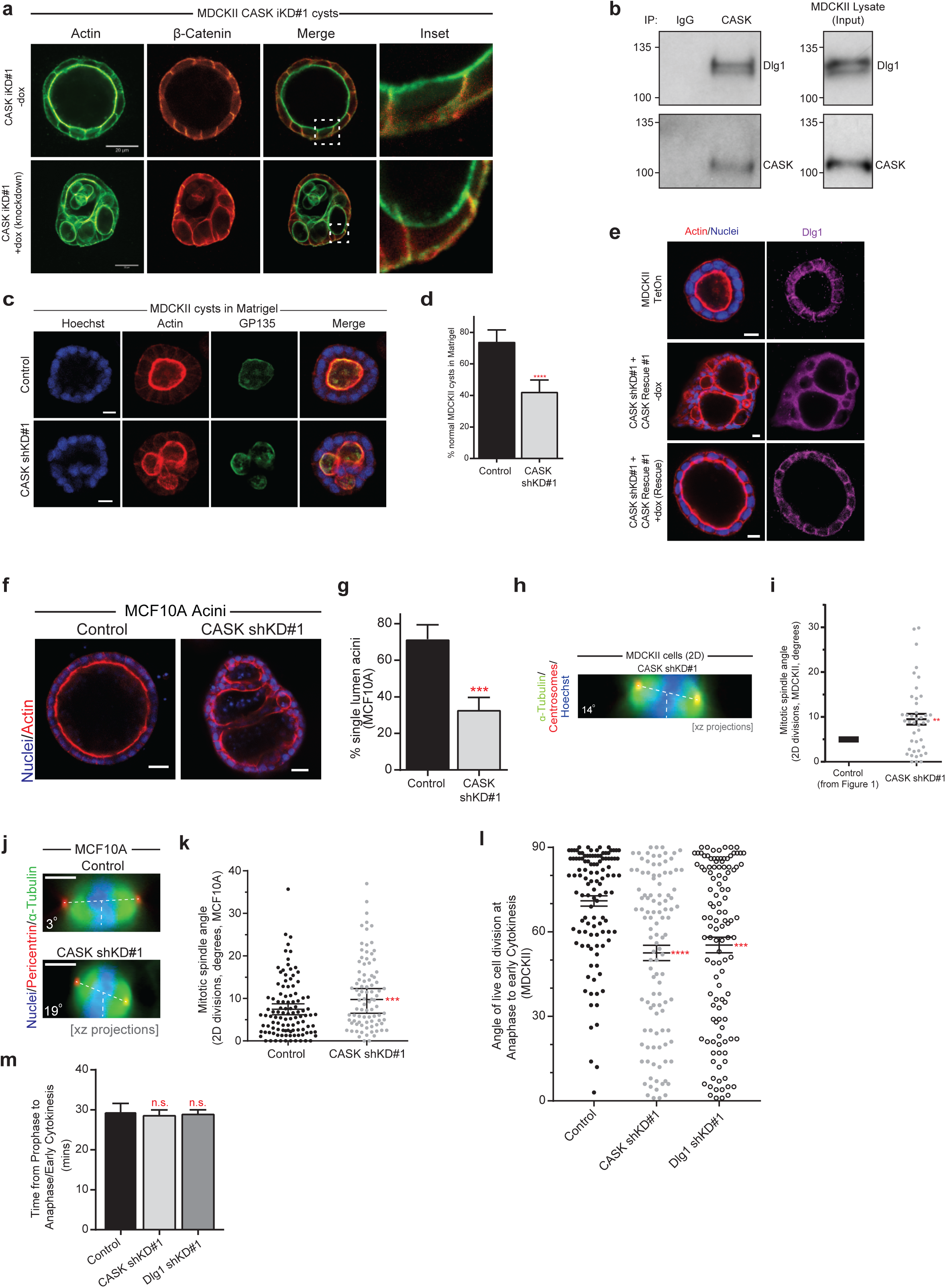
a) MDCKII cysts showing strong basolateral staining of β-Catenin in both Control (-dox) and CASK iKD#1 (+dox). b) Western blot for CASK and Dlg1, following CASK immunoprecipitation. c) Confocal images of cysts from MDCKII cells transfected with either Control (Non-Targeting) or CASK shRNA grown in 2% Matrigel and displaying either a normal (top panels) or a multilumen phenotype (bottom panels). d) Quantification of cysts with normal lumens as depicted in (c), N=3 independent experiments, more than 100 cysts examined per experiment; **** p=2.6×10^−5^ (unpaired Student’s T-test, 2 tailed). e) Confocal images of MDCKII cysts showing basolateral localisation of Dlg1 in Control cysts (MDCKII Tet-On, top panel), loss of Dlg1 lateral staining following CASK knockdown (-dox, middle panel) and restoration of basolateral Dlg1 staining upon CASK re-expression (+dox, bottom panel). f) Confocal images of MCF10A accini, grown on Matrigel, showing multi-acini structures following transfection with CASK shKD#1 to knockdown CASK, but not following transfection with the Non-Targeting shRNA (Control). g) Quantification of normal MCF10A accini from experiments as in (f). N=4 independent experiments for both conditions, with more than 100 acini examined per experiment; *** p=0.0003 (unpaired Student’s T-test, 2 tailed). Values are mean +/- SEM. h) Representative images of XZ projections from MDCKII cells grown in 2D, showing tilted mitotic spindles at metaphase after CASK knockdown, and annotated to show guide lines and spindle angles. i) Quantification of spindle angles from MDCKII Control (Non-Targeting) and CASK knockdown cells grown in 2D, n=42 from three independent experiments; ** p= 0.0046 (Mann-Whitney test), compared with control (control mean and SEM are shown as in Supplementary Figure 1b). Error bars show mean +/- SEM. Data analysed using Kruskal-Wallis test. j) XZ projection of metaphase MCF10A cells showing tilting of the metaphase spindle after knockdown of CASK (bottom panel). Scale bar is 5 µm. k) Quantification of 2D spindle angle in MCF10A cells from experiments as in (j). n=107/88 cells pooled from three independent experiments; *** p=0.00052 (Mann-Whitney test). Error bars show mean +/- SEM. l) Quantification of cell division angles from live imaging experiments, only from cells in anaphase, telophase or early cytokinesis (n=112/110/116 cells from three independent replicates); **** p=2.9×10^−6^, *** p=0.00031. Values are mean +/- SEM (Kruskal-Wallis test.) m) In cases where individual cell divisions could be followed from prophase through to anaphase/telophase, the timing of progression was recorded (interval between images was 7.5 minutes). Graph shows mean, error bars show 1SD (n=51/30/24 from three biological replicates). n.s. = not significant (p= 0.9134 and p= 0.9653 for CASK and Dlg1 shRNA respectively). (One-way Anova with multiple comparisons.)

**Supplementary Figure 3.**
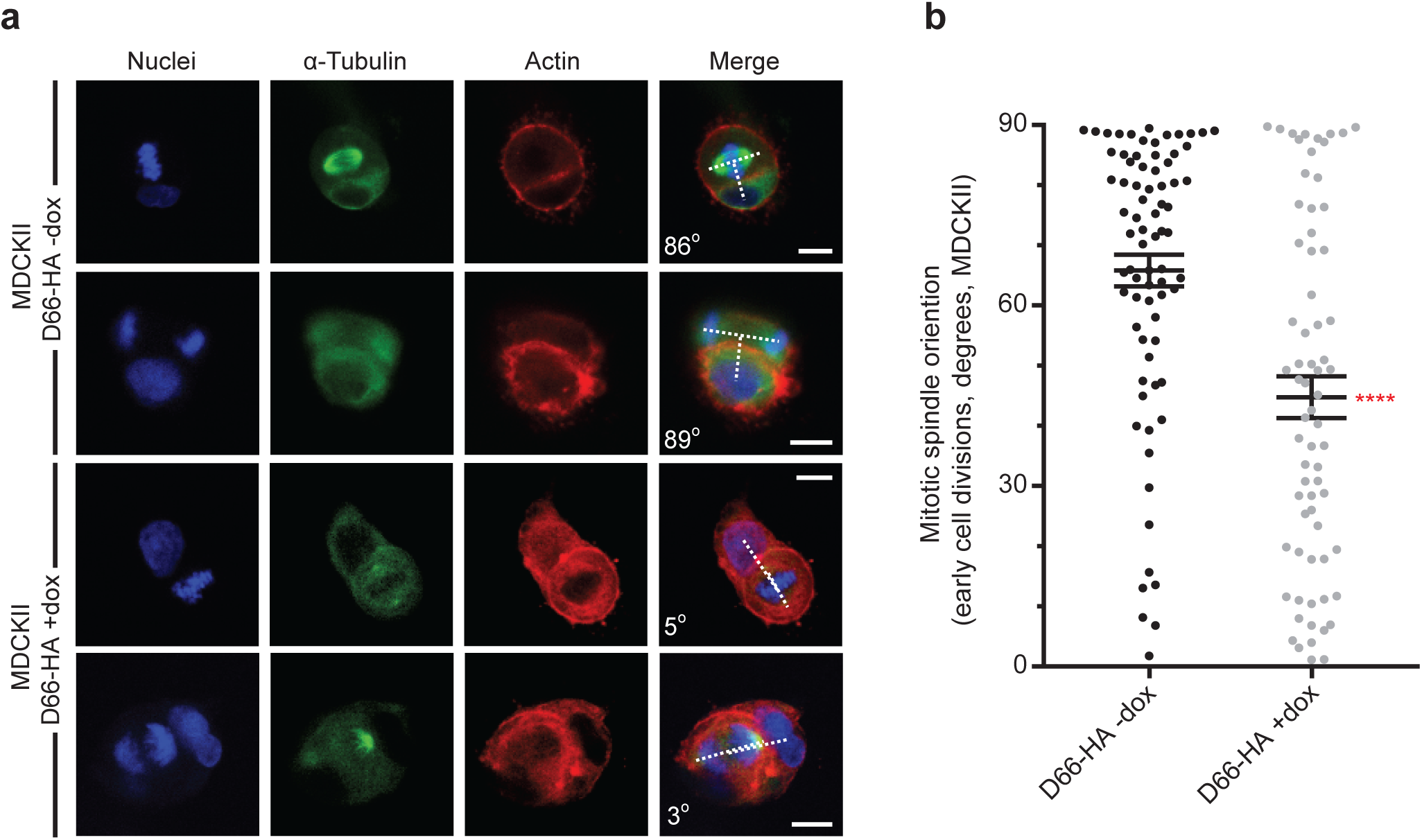
a) Example images of early cell divisions in MDCKII cysts showing misoriented divisions following expression of D66-HA. b) Quantification of spindle orientation angles in early MDCKII cysts with and without D66-HA expression, n=76/69 from three independent experiments; **** p=2.3×10^−5^ (Mann Whitney Test). Error bars show mean +/- SEM.

**Supplementary Figure 4.**
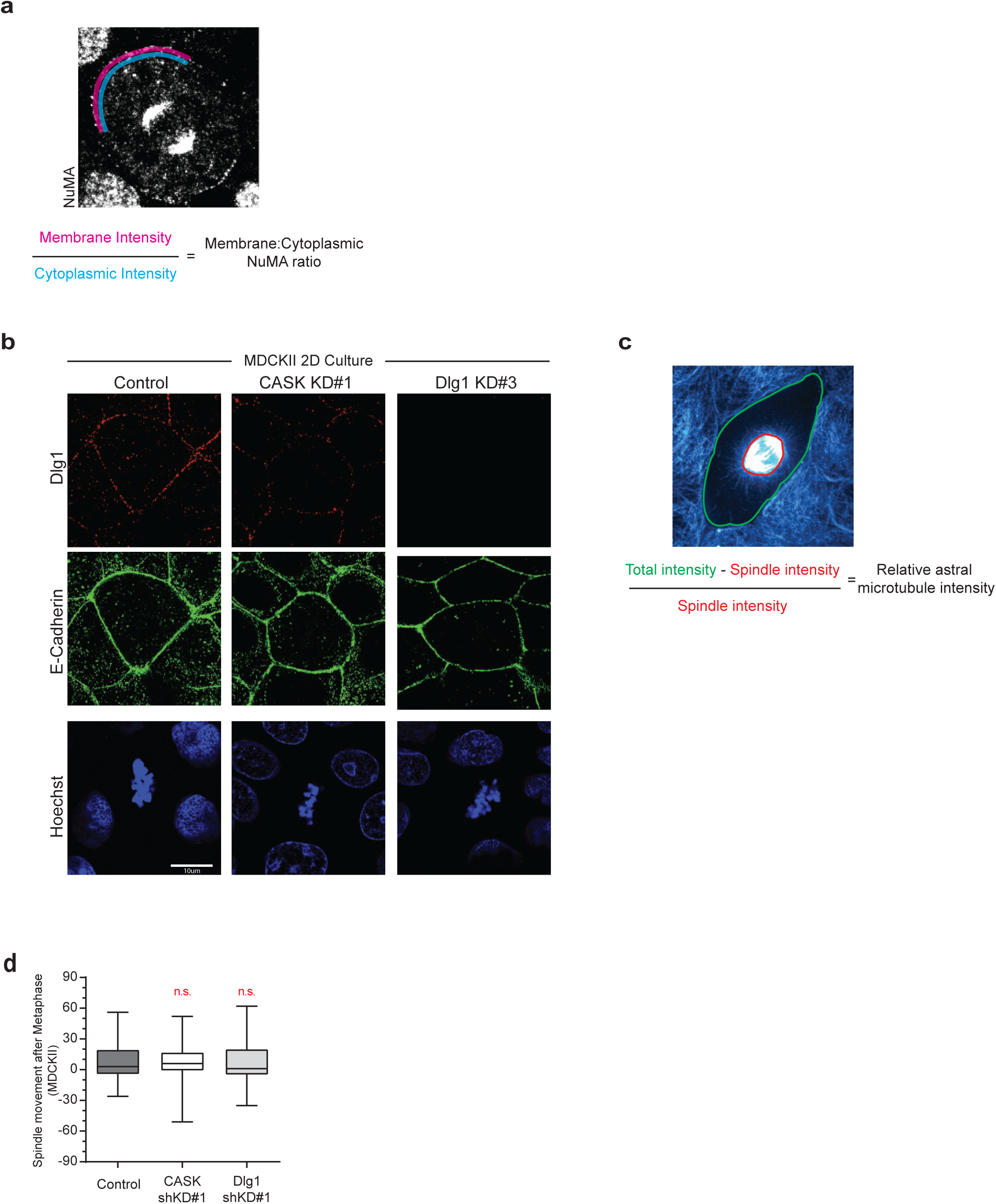
a) Schematic of Membrane:Cytoplasmic ratio quantification used for LGN and NuMA imaging; purple line represents Membrane intensity quantification, blue line represents Cytoplasmic intensity quantification. b) Single plane confocal images of MDCKII cells plated in 2D, showing normal E-Cadherin staining at cell junctions which is maintained following CASK and Dlg1 knockdown. c) Schematic of relative astral microtubule intensity quantification; red line indicates area of the mitotic spindle, green line indicates total cell area. d) Box and whisker plot showing change in mitotic angle between metaphase and anaphase to early cytokinesis for individual cells where the angle at both stages could be measured (n=93/99/103 from three biological replicates). Box shows 25-75 percentile marked with the median, whiskers show min/max. n.s. = not significant (p=0.93 and 0.95 for CASK and Dlg1 shRNA respectively).

### Supplementary Movie Legends

**Supplementary Movie 1**

Non-Targeting shRNA-expressing control cyst, grown on Matrigel, showing normally oriented cell divisions; cell division starting at 0 minutes corresponds to division in Figure 3a. Other divisions in the movie are also oriented in the plane of the epithelium. Left panel – merged image. Middle panel

– Actin Chromobody, tagRFP. Right panel – Histone-2B-CFP. 4 frames/second.

**Supplementary Movie 2**

CASK shKD#1 cyst, grown on Matrigel, showing a number of abnormally oriented cell divisions; cell division starting at 0 minutes corresponds to division in Figure 3b. Other divisions in the movie are also abnormally oriented, and what appears to be a new lumen forms towards the end of the

recording. Left panel – merged image. Middle panel – Actin Chromobody, tagRFP. Right panel – Histone-2B-CFP. 4 frames/second.

**Supplementary Movie** 3

Dlg1 shKD#1 cyst, grown on Matrigel, showing an abnormally oriented cell division starting at 0 minutes (corresponds to division in Figure 3c). Left panel – merged image. Middle panel – Actin Chromobody, tagRFP. Right panel – Histone-2B-CFP. 4 frames/second.

## References

1. di Pietro, F., A. Echard, and X. Morin, Regulation of mitotic spindle orientation: an integrated view. EMBO Rep, 2016. 17(8): p. 1106–30.

2. Bergstralh, D.T., T. Haack, and D. St Johnston, Epithelial polarity and spindle orientation: intersecting pathways. Philos Trans R Soc Lond B Biol Sci, 2013. 368(1629): p. 20130291.

3. Pease, J.C. and J.S. Tirnauer, Mitotic spindle misorientation in cancer--out of alignment and into the fire. J Cell Sci, 2011. 124(Pt 7): p. 1007–16.

4. Lu, M.S. and C.A. Johnston, Molecular pathways regulating mitotic spindle orientation in animal cells. Development, 2013. 140(9): p. 1843–56.

5. Du, Q., et al., LGN blocks the ability of NuMA to bind and stabilize microtubules. A mechanism for mitotic spindle assembly regulation. Curr Biol, 2002. 12(22): p. 1928–33.

6. Saadaoui, M., et al., Dlg1 controls planar spindle orientation in the neuroepithelium through direct interaction with LGN. J Cell Biol, 2014. 206(6): p. 707–17.

7. Bergstralh, D.T., H.E. Lovegrove, and D. St Johnston, Discs large links spindle orientation to apical-basal polarity in Drosophila epithelia. Curr Biol, 2013. 23(17): p. 1707–12.

8. Gloerich, M., et al., Cell division orientation is coupled to cell-cell adhesion by the E-cadherin/LGN complex. Nat Commun, 2017. 8: p. 13996.

9. Bergstralh, D.T., N.S. Dawney, and D. St Johnston, Spindle orientation: a question of complex positioning. Development, 2017. 144(7): p. 1137–1145.

10. Hao, Y., et al., Par3 controls epithelial spindle orientation by aPKC-mediated phosphorylation of apical Pins. Curr Biol, 2010. 20(20): p. 1809–18.

11. Zheng, Z., et al., LGN regulates mitotic spindle orientation during epithelial morphogenesis. J Cell Biol, 2010. 189(2): p. 275–88.

12. Humbert, P.O., et al., Control of tumourigenesis by the Scribble/Dlg/Lgl polarity module. Oncogene, 2008. 27(55): p. 6888–907.

13. Firestein, B.L. and C. Rongo, DLG-1 is a MAGUK similar to SAP97 and is required for adherens junction formation. Mol Biol Cell, 2001. 12(11): p. 3465–75.

14. Lee, S., et al., A novel and conserved protein-protein interaction domain of mammalian Lin-2/CASK binds and recruits SAP97 to the lateral surface of epithelia. Mol Cell Biol, 2002. 22(6): p. 1778–91.

15. Lozovatsky, L., et al., CASK deletion in intestinal epithelia causes mislocalization of LIN7C and the DLG1/Scrib polarity complex without affecting cell polarity. Mol Biol Cell, 2009. 20(21): p. 4489–99.

16. Mukherjee, K., et al., CASK Functions as a Mg2+-independent neurexin kinase. Cell, 2008. 133(2): p. 328–39.

17. Atasoy, D., et al., Deletion of CASK in mice is lethal and impairs synaptic function. Proc Natl Acad Sci U S A, 2007. 104(7): p. 2525–30.

18. Hackett, A., et al., CASK mutations are frequent in males and cause X-linked nystagmus and variable XLMR phenotypes. Eur J Hum Genet, 2010. 18(5): p. 544–52.

19. Hsueh, Y.P., The role of the MAGUK protein CASK in neural development and synaptic function. Curr Med Chem, 2006. 13(16): p. 1915–27.

20. Lin, E.I., O. Jeyifous, and W.N. Green, CASK regulates SAP97 conformation and its interactions with AMPA and NMDA receptors. J Neurosci, 2013. 33(29): p. 12067–76.

21. Wang, T.F., et al., Identification of Tbr-1/CASK complex target genes in neurons. J Neurochem, 2004. 91(6): p. 1483–92.

22. Mack, N.A., et al., beta2-syntrophin and Par-3 promote an apicobasal Rac activity gradient at cell-cell junctions by differentially regulating Tiam1 activity. Nat Cell Biol, 2012. 14(11): p. 1169–80.

23. Seldin, L. and I. Macara, Epithelial spindle orientation diversities and uncertainties: recent developments and lingering questions. F1000Res, 2017. 6: p. 984.

24. Hart, K.C., et al., E-cadherin and LGN align epithelial cell divisions with tissue tension independently of cell shape. Proc Natl Acad Sci U S A, 2017. 114(29): p. E5845–E5853.

25. Wang, X., et al., E-cadherin bridges cell polarity and spindle orientation to ensure prostate epithelial integrity and prevent carcinogenesis in vivo. PLoS Genet, 2018. 14(8): p. e1007609.

26. Du, Q. and I.G. Macara, Mammalian Pins is a conformational switch that links NuMA to heterotrimeric G proteins. Cell, 2004. 119(4): p. 503–16.

27. Peyre, E., et al., A lateral belt of cortical LGN and NuMA guides mitotic spindle movements and planar division in neuroepithelial cells. J Cell Biol, 2011. 193(1): p. 141–54.

28. Seldin, L., A. Muroyama, and T. Lechler, NuMA-microtubule interactions are critical for spindle orientation and the morphogenesis of diverse epidermal structures. Elife, 2016. 5.

29. Cohen, A.R., et al., Human CASK/LIN-2 binds syndecan-2 and protein 4.1 and localizes to the basolateral membrane of epithelial cells. J Cell Biol, 1998. 142(1): p. 129–38.

30. Bosveld, F., et al., Epithelial tricellular junctions act as interphase cell shape sensors to orient mitosis. Nature, 2016. 530(7591): p. 495–8.

31. Saadaoui, M., et al., Loss of the canonical spindle orientation function in the Pins/LGN homolog AGS3. EMBO Rep, 2017. 18(9): p. 1509–1520.

32. Awad, A., et al., SHIP2 regulates epithelial cell polarity through its lipid product, which binds to Dlg1, a pathway subverted by hepatitis C virus core protein. Mol Biol Cell, 2013. 24(14): p. 2171–85.

33. Lancaster, M.A. and J.A. Knoblich, Spindle orientation in mammalian cerebral cortical development. Curr Opin Neurobiol, 2012. 22(5): p. 737–46.

34. Yang, Y., et al., CYLD regulates spindle orientation by stabilizing astral microtubules and promoting dishevelled-NuMA-dynein/dynactin complex formation. Proc Natl Acad Sci U S A, 2014. 111(6): p. 2158–63.

35. Debnath, J. and J.S. Brugge, Modelling glandular epithelial cancers in three-dimensional cultures. Nat Rev Cancer, 2005. 5(9): p. 675–88.

36. Naim, E., et al., Mutagenesis of the epithelial polarity gene, discs large 1, perturbs nephrogenesis in the developing mouse kidney. Kidney Int, 2005. 68(3): p. 955–65.

37. Bergstralh, D.T., H.E. Lovegrove, and D. St Johnston, Lateral adhesion drives reintegration of misplaced cells into epithelial monolayers. Nat Cell Biol, 2015. 17(11): p. 1497–1503.

38. Woodcock, S.A., et al., Tiam1-Rac signaling counteracts Eg5 during bipolar spindle assembly to facilitate chromosome congression. Curr Biol, 2010. 20(7): p. 669–75.

39. Whalley, H.J., et al., Cdk1 phosphorylates the Rac activator Tiam1 to activate centrosomal Pak and promote mitotic spindle formation. Nat Commun, 2015. 6: p. 7437.

